# Empirical Geometry-Guided Modeling Enables Robust, High-Throughput Collagen Structure Prediction

**DOI:** 10.64898/2026.09.05.749573

**Authors:** Bruno Howard, Maxwell Bauer, Markus J. Buehler

## Abstract

Protein structure prediction is increasingly dominated by large learned generative models, yet for proteins governed by strong structural constraints, much of the relevant conformational space may be captured by substantially more compact representations. Collagen provides a compelling test case: its repeating Gly-X-Y sequence, restricted backbone conformations, and conserved triple-helical topology define a highly constrained structural manifold. Here, we develop a Collagen-specific Deterministic Structure Modeler (CDSM), extending the empirical geometric parameterization of THeBuScr into a robust, all-atom structure-prediction pipeline, and benchmark it against AlphaFold 3, Boltz-2, Chai-1, and Protenix-v1 on 80 experimentally resolved collagen triple-helical structures. CDSM increases coverage of this benchmark from 8.8% for native THeBuScr to 93.8%. Across the 75 structures successfully predicted by all methods, CDSM closely reproduces experimental backbone and global geometry, with fewer large-error predictions, while the learned models achieve modestly higher local and side-chain accuracy. When evaluated only on structures deposited after each learned model’s training-data cutoff, CDSM becomes more competitive, with aggregate win rates increasing from 29% to 55% for TM-score and from 40% to 65% for backbone RMSD, while CDSM performance remains comparatively stable across the same subsets. CDSM generates structures in 2.4 s on a single CPU core at approximately $3 × 10^−5^ per structure, making it 400 to 790× cheaper than the learned methods even when each is run on its lowest-cost compatible GPU. These results demonstrate how identifying and explicitly encoding relevant geometric constraints can dramatically compress a structure-prediction problem, retaining much of the accuracy at orders-of-magnitude lower computational cost. More broadly, CDSM illustrates the potential for compact, executable scientific representations that complement large learned models, and motivates future AI-driven searches over algorithms, representations, and empirical parameterizations to discover and continually refine such models for other structural protein domains.

## 1 INTRODUCTION

Collagen is the most abundant protein in the human body, comprising approximately one third of total protein mass [1,2]. It provides the mechanical framework for connective tissues such as skin, bone, and tendon, plays critical roles in cellular repair, angiogenesis and tissue development, and is implicated in a range of pathological conditions [3–5]. Its defining structural feature is a repeating Gly-X-Y amino acid motif across three left-handed polypeptide α-chains, which self-assemble into a characteristic right-handed triple helix with a one-residue stagger between adjacent α-chains [6]. Variations at the X and Y positions, which are commonly occupied by proline (Pro) and 4-hydroxyproline (Hyp), respectively (together comprising over 20% of collagen’s amino acid content) [7], tune the protein’s stability and mechanical behavior, with hydroxyproline in particular playing an important role in stabilizing the triple helix [8, 9].

The protein’s physiological importance and structural distinctiveness have motivated numerous high-resolution X-ray crystallography studies [10–12]. These studies have advanced our understanding of the triple-helix structure and underpin more recent efforts toward rational collagen design for applications ranging from tissue scaffolds to wound-healing matrices [13–15]. Indeed, as computational capabilities evolve, collagen is becoming a natural target for inverse design: tailoring sequence-level features to achieve desired functional properties. Recent work has shown that large language models trained on only 633 collagen mimetic peptide (CMP) sequence-to-T_m_ pairs can predict melting temperature (a proxy for protein stability) with high accuracy (R^2^ = 0.84), demonstrating that sequence-property relationships can be learned from sequence alone [1, 16]. However, collagen performs its remarkable biological function as a three-dimensional structure, and its mechanical and biophysical properties arise from unique geometric features and intermolecular interactions that are not explicitly represented in a single chain sequence. Structure-aware approaches therefore offer the potential to capture richer information relevant to collagen function and to enable more physically interpretable computational models [17, 18].

However, such models require large amounts of structural data. Although collagen is among the best structurally characterized fibrous proteins, experimentally determined structural diversity remains limited [19, 20]. Since the first triple-helical peptide crystal structure was reported in 1994 [10], only a few hundred collagen triple-helical structures have been experimentally resolved, compared with ~230,000 predominantly globular protein structures in the PDB (RCSB PDB, 06/26). These structures have provided critical insight into collagen architecture, but remain insufficient for large-scale computational studies. Consequently, accurate generation of collagen structures from sequence is an important enabling step.

Several approaches exist for computational structure prediction. Molecular dynamics (MD) can resolve atomic-level detail through physics-based simulation but is computationally expensive, with exhaustive conformational sampling generally prohibitive at the scale required for design screening [21, 22]. More recently, deep-learning models such as AlphaFold (Google DeepMind) have transformed protein structure prediction [23], with newer approaches building on similar architectures to extend these capabilities [24–27]. However, their performance on repetitive fibrous proteins such as collagen remains less established than on globular proteins, particularly given the structural-data imbalance already described. Collagen also depends critically on post-translational hydroxylation of proline [28], introducing complexity beyond prediction based on the conventional 20 amino acids. Such challenges persist even for recent models such as AlphaFold 3, which has shown mixed performance on collagen-domain assemblies [29, 30].

An alternative approach is to exploit collagen’s unusually strong structural constraints directly. Its repetitive primary sequence, restricted backbone conformations, and conserved triple-helical topology make collagen particularly well suited to a compact empirical geometric representation [28, 31]. Rainey & Goh (2002) derived a statistical parameter set from available crystal structures encoding backbone dihedral angles and local helical geometry, enabling prediction of backbone and Cβ atom positions directly from sequence [20], and packaged it as the Triple Helical collagen Builder Script, or *THeBuScr* [32]. Here, we extend this approach into the *Collagen-specific Deterministic Structure Modeler* (CDSM), a high-throughput pipeline for generating complete all-atom collagen structures directly from sequence. CDSM retains *THeBuScr’s* parameterization as its structural foundation, while adding sequence reframing, terminal reconstruction, and chain-register correction to accommodate more diverse collagen sequences, including unequal chain lengths and incomplete Gly-X-Y triplets. The resulting backbone is completed with side-chain atoms using *AMBER tleap* and subjected to restrained energy minimization, producing full all-atom structures while preserving the underlying parameterized triple-helical geometry.

Despite advances in AI-based protein structure prediction, no collagen-specific benchmark has systematically compared these methods. Here, we evaluate AlphaFold 3, Chai-1, Protenix-v1 and Boltz-2 alongside our proposed CDSM pipeline using experimentally resolved collagen triple helices as a benchmark. We assess structural accuracy and computational cost to evaluate suitability for high-throughput collagen structure generation. More broadly, this comparison tests whether explicitly encoding a protein’s underlying geometric constraints can provide a compact yet accurate representation of its structural manifold. Emerging self-revising AI systems may provide a means to autonomously search over algorithms and empirical parameterizations to discover such compact, executable representations, and to continually reparametrize and refine them as new data become available [33].

## 2 MATERIALS AND METHODS

This study benchmarks our proposed Collagen-specific Deterministic Structure Modeler (CDSM) against four frontier deep-learning structure predictors - AlphaFold 3, Chai-1, Protenix-v1, and Boltz-2 [24–27] - using experimentally resolved collagen triple helices as structural references. The study aims to assess the relative suitability of these methods for high-throughput collagen structure prediction and how closely a compact, geometric representation can approach the performance of general-purpose learned models. This section describes the benchmark dataset, structure prediction methods, and evaluation procedures.

### 2.1 Dataset

To evaluate collagen structure prediction methods, we curate a set of experimentally resolved triple-helix structures with their corresponding crystallographically observed sequences. The benchmark dataset comprises 80 collagen triple-helix structures (30 homotrimers, 50 heterotrimers), which serve as ground truth. Structures were identified via the *RCSB Search API (v2)* using the keywords “collagen” or “triple helix”, restricted to protein-only entries with deposited polymer-chain instances that are positive multiples of three only. Allowing multiples of three ensures asymmetric units containing more than one crystallographically independent copy of a triple helix are included. The search identified 758 candidate structures at the time of retrieval. This was reduced to 80 structures through a sequential filtering process.

Candidate structures were retained only when all observed residues belonged to an alphabet comprising the 20 canonical amino acids and (4R)-4-hydroxyproline (HYP). To distinguish collagen triple helices from unrelated trimeric proteins captured by the broad text search, structures were required to have an observed glycine fraction of at least 25%, consistent with the approximately one-in-three glycine periodicity of canonical collagen. Structures were additionally excluded on the basis of structural completeness (residue-numbering gap >5), chain length (10-200 residues), and exact-sequence deduplication. For asymmetric units containing multiple helices, chains were grouped into triple helices by spatial distance and one helix was retained. The resulting set of 80 sequences ranges from 16 to 36 residues per chain (median 27, mean 26.8) and spans 85 unique Gly-X-Y triplets, where Proline and Hydroxyproline account for approximately 80% of all non-glycine residues.

To our knowledge, this dataset represents the most comprehensive benchmark set of canonical collagen triple-helix structures supported by all methods evaluated here. The complete curation procedure from search query, to rejection counts, PDB identifiers, experimental resolutions, sequences, and structure-level metadata are provided in the *Supplementary Information, S1*.

### 2.2 Methods: Structure Prediction

Each set of three deposited chain sequences is used to generate predicted collagen structures using (1) our collagen-specific deterministic structure modeler (CDSM) and (2) the four deep-learning structure prediction models, AlphaFold 3, Boltz-2, Chai-1, and Protenix-v1. For each, the input is restricted to amino acid sequence only, and structures are generated without reference or access to the experimental coordinates.

#### 2.2.1 Collagen-specific Deterministic Structure Modeler (CDSM)

Our proposed *Collagen-specific Deterministic Structure Modeler* is built on the Triple Helical collagen Builder Script (THeBuScr) [32]. THeBuScr constructs an idealized collagen triple helix backbone assuming glycine at every third residue and a sequence capable of forming a stable triple helix. Backbone coordinates (N, Cα, C, O) are generated using a statistical parameter set derived from crystallographic measur-e ments of collagen-like peptides [20]. Sequence-dependent helical propensity estimates from Persikov et al. [34] are used to distinguish different conformational regions. A weighted sliding window over five neighboring residues assigns each glycine-leading triplet to an imino-acid-rich (IR) or amino-acid-rich (AR) conformational state, which, together with each residue’s position in the Gly-X-Y repeat, determines its local helical geometry. The three chains are arranged into the characteristic staggered triple helix using statistically derived chain-to-chain axial translations [20]. Side-chain geometry is not modeled.

THeBuScr is a remarkably successful idealized collagen backbone builder, but has several limitations. It assumes that glycine occupies every third residue position, that each chain begins with glycine, that all three chains comprise complete triplets of equal length, and it reconstructs only backbone and Cβ atoms. Consequently, it cannot accommodate incomplete terminal triplets, differing chain lengths within homo- or heterotrimers, or generate complete all-atom structures.

CDSM addresses these limitations through eight steps across five stages (Fig. 3), enabling full-length all-atom structures from sequences with arbitrary chain starting positions, incomplete terminal triplets, and unequal or overhanging chain lengths. These capabilities are achieved through sequence reframing, terminal reconstruction by helical extension or geometric extrapolation, chain-register correction, side-chain construction, and restrained energy minimization. The resulting Python-native, high-throughput pipeline generates complete collagen structures directly from sequence without model training.

**Figure 1:**
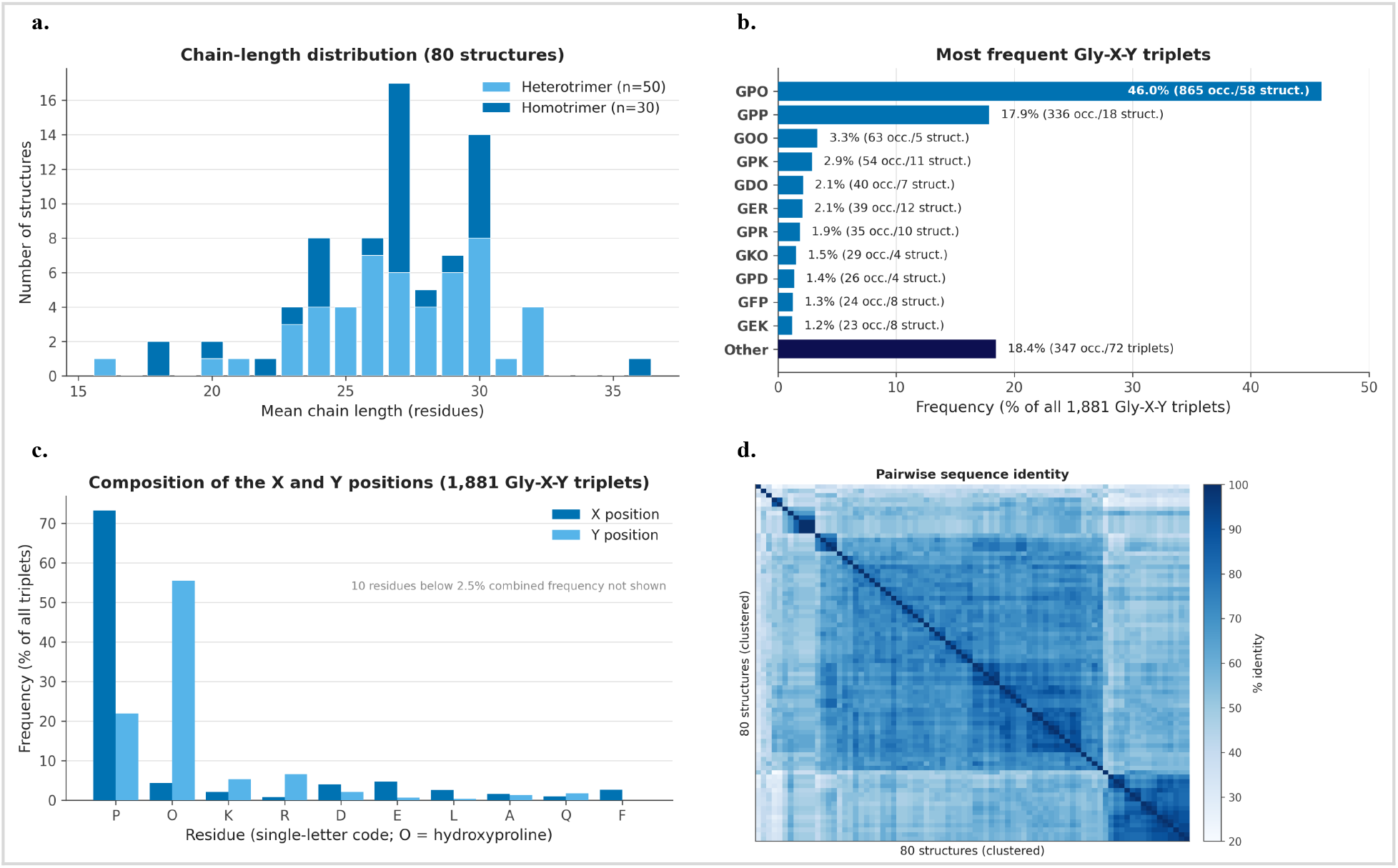
Dataset. **a**. Mean chain length (average of three chains per structure). Sequences range from 16 to 36 residues per chain (median 27, mean 26.8). **b**. The dataset spans 85 unique Gly-X-Y triplets, of which 44 occur only once, in 44 unique structures. Gly-Pro-Hyp (GPO) is the dominant motif, representing 46% of all triplets; **c**. Pro and Hyp together account for approximately 80% of all non-glycine residues. Pro dominates the X position (~75%) while Hyp dominates Y (~56%), reflecting the known enzymatic preference for hydroxylation at the Y position. **d**. Pairwise sequence identity, calculated across each structure’s chain A, reveals a median 61% sequence similarity, reflecting the highly repetitive structure of collagen (where each third glycine occupying position shares practically 100% similarity). However, only 7% of sequences exceed 80% similarity, reflecting significant diversity amongst the non-repeating parts of the Gly-X-Y scaffold.

**Figure 2:**
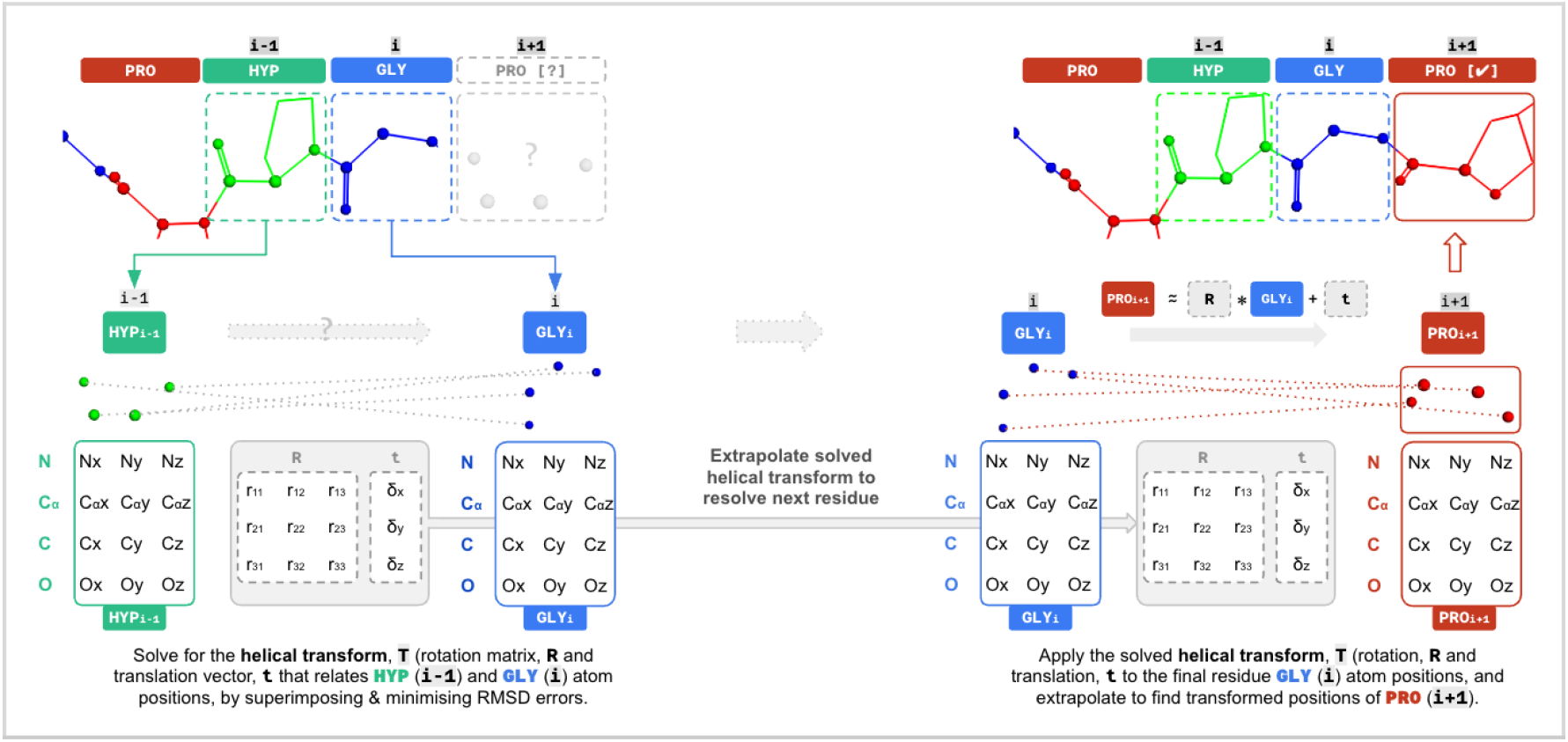
Illustration of the Kabsch extrapolation to resolve incomplete triplet terminal residue backbone atom coordinates (N, Cα, C, O). The Kabsch algorithm seeks to resolve the helical transform consisting of a rotation matrix and a translation vector by minimizing the RMSD error between the two known residue atom coordinates. Here, the algorithm is used to determine the transformation (R,t) from HYP (i-1) to GLY (i). The same transformation is then extrapolated and applied to GLY (i) to approximate the backbone atom positions of PRO (i+1).

**Figure 3:**
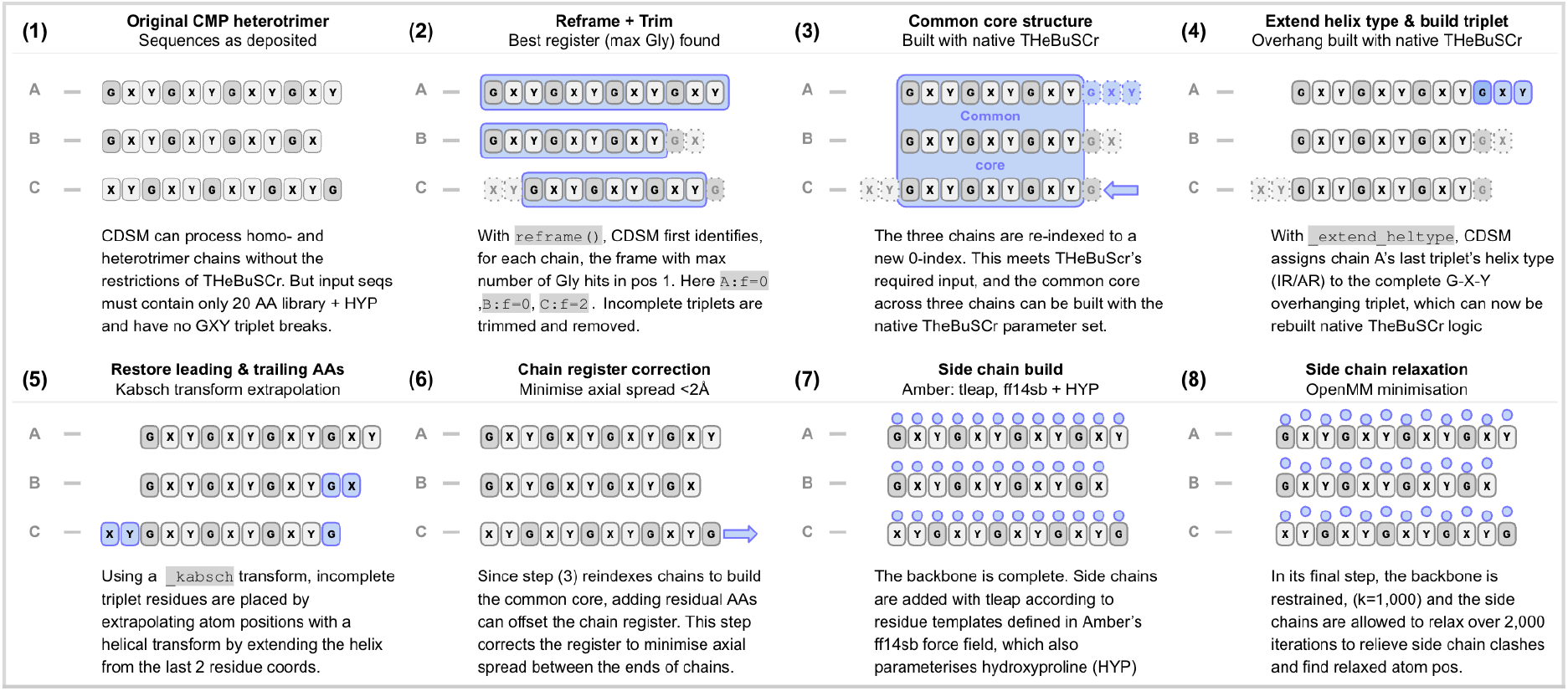
Collagen-specific Deterministic Structure Modeler (CDSM) pipeline. CDSM extends THeBuScr to accommodate non-canonical chain termini and unequal chain lengths through sequence reframing and trimming, common-core construction, terminal reconstruction, and chain-register correction, followed by all-atom side-chain construction and restrained energy minimization.

**Figure 4:**
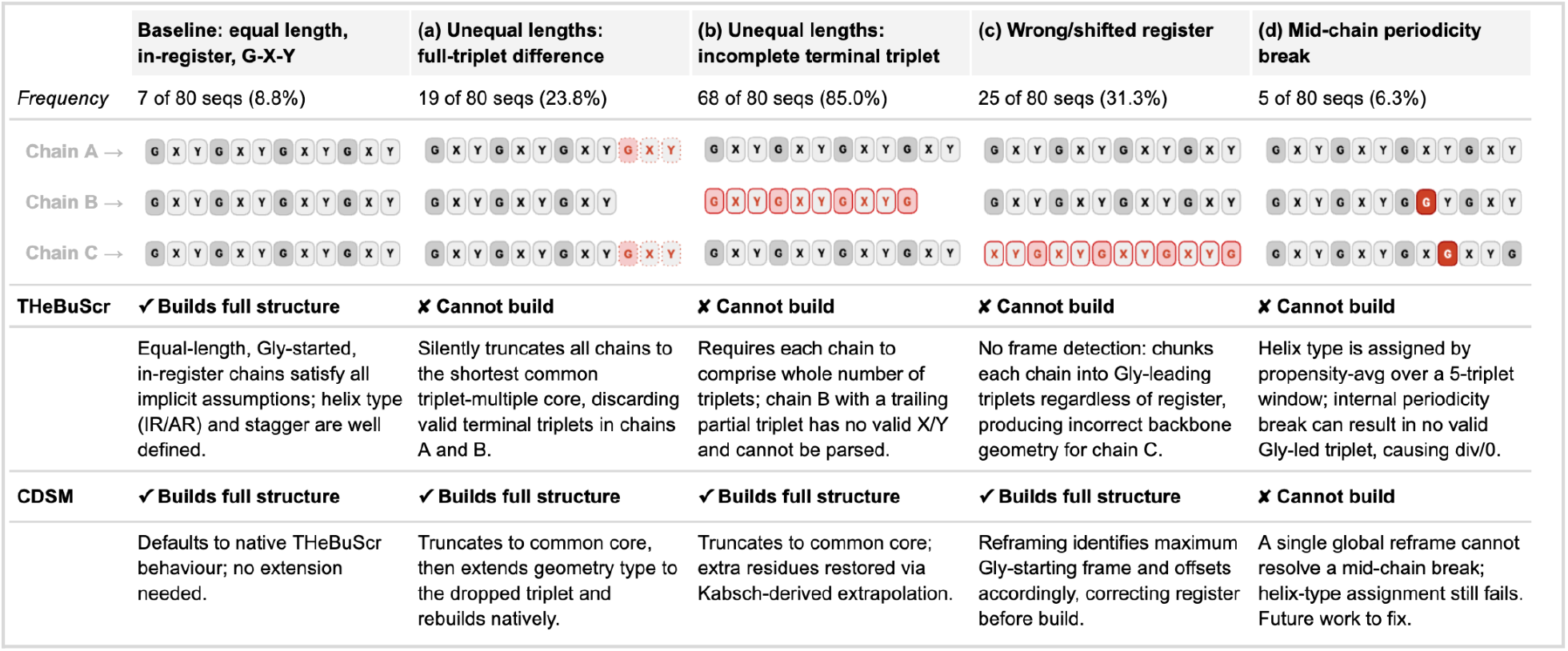
CDSM extends THeBuScr to accommodate diverse collagen chain geometries. Representative sequences illustrate the limitations of native THeBuScr and their resolution by CDSM. CDSM resolves unequal chain lengths, incomplete terminal triplets, and shifted Gly-X-Y register, increasing benchmark coverage from 7/80 (8.8%) to 75/80 (93.8%). Internal disruptions of Gly-X-Y periodicity remain unsupported.

Stage 1: Sequence reframing and common-core construction *(Fig. 3.1-3)* - THeBuScr applies a fixed geometry to successive Gly-X-Y triplets but assumes that the first residue position of the repeat is occupied by a glycine, resulting in X- or Y-starting chains to be built incorrectly. To address this, each chain is first reframed so that the maximum number of glycines occupy the first position of Gly-X-Y triplets across the length of each chain. This causes up to two residues from incomplete triplets to be removed from the N- and C-termini, together with any whole triplets extending beyond the common triple-helical core shared by all three chains. This preprocessing step maximizes the number of triplets within the core of the structure whose atom locations can be determined according to the Rainey-Goh parameter set. The three chains are then zero-indexed to establish the canonical one-residue axial stagger of the collagen triple helix.

Stage 2: Terminal reconstruction *(Fig. 3.4-5)* - Once the common core is constructed, trimmed regions are reconstructed with two different mechanisms. Whole-triplet overhangs in chains longer than the common core are reconstructed by propagating the terminal amino- or imino-type helical class to the overhanging triplet, allowing backbone to be extended using THeBuScr’s native parameterization. In contrast, incomplete terminal triplets cannot be assigned a local helical propensity and are reconstructed by geometrical extrapolation: a rigid-body screw transformation is derived from the backbone coordinates of the two built terminal residues using Kabsch superposition and is applied forward at the C-terminus, or in inverse at the N-terminus, to restore trimmed residue coordinates. Together, these steps restore the complete experimental sequence rather than restricting construction to an equal-length central core.

Stage 3: Chain-register correction *(Fig. 3.6)* - Following reframing, terminal reconstruction, and application of the canonical one-residue stagger, chains of unequal length may remain axially misregistered, producing splayed termini not observed experimentally. The third stage corrects the structure’s chain register by evaluating integer whole-triplet shifts for each chain, and selecting the threading that minimizes terminal axial spread. A shift is applied only if it reduces terminal overhang by ≥2 Å. Shifts are restricted to complete triplets so that the glycine register and identities of the X and Y positions remain unchanged.

Stage 4: Side-chain construction *(Fig. 3.7)* - Once the backbone is built, side-chain atom coordinates are determined with *AmberTools’ tleap* [35] with the *ff14SB* force field [36], including explicit parameters for hydroxyproline (HYP), the predominant post-translational modification in collagen. Chain identifiers are restored using ParmEd [37], and hydrogen atoms are removed for structural evaluation.

Stage 5: Restrained energy minimization (Fig. 3.8) - The completed all-atom structures are energy-minimized using OpenMM 8.4 [38] with the OBC2 implicit solvent model [39]. Harmonic positional restraints (1000 kJ mol^−1^ nm^−2^) are applied to the backbone to preserve the triple-helical geometry while allowing side chains and local steric clashes to relax. Minimization is performed to default force tolerance with a maximum of 2,000 iterations.

While CDSM substantially extends THeBuScr, it retains one fundamental limitation: extended internal disruptions of Gly-X-Y periodicity cannot be constructed. When no canonical triplets remain within the propensity-averaging window, the local propensity calculation becomes undefined, preventing assignment of the helix conformation required for backbone construction. As a result, CDSM generates complete all-atom structures for 75 of the 80 benchmark targets (compared with 7 of 80 using native THeBuScr), with the remaining five failures caused by extended Gly-X-Y disruptions. Ongoing work aims to address this limitation through alternative treatments of local helical propensities across interrupted motifs.

#### 2.2.2 Deep-Learning Structure Prediction Models: Boltz-2, Chai-1, Protenix-v1, AlphaFold 3

All four learned models were provided with identical biological specifications: three-chain assemblies with stoichiometry matching the experimental structure (homotrimer = one sequence entity with copy count 3; heterotrimer = three distinct entities), and hydroxyproline encoded explicitly via the PDB Chemical Component Dictionary code HYP rather than being collapsed to proline. HYP encoding position indexing conventions differ between models (0-indexed for Boltz-2; inline (HYP) notation for Chai-1; 1-indexed ptmPosition for Protenix-v1 and AlphaFold 3) and were validated prior to batch runs.

A comparison of MSA-based and single-sequence predictions on five representative structures showed no consistent benefit from including MSA information. All models were therefore run in single-sequence mode, with other inference settings kept equivalent where supported. Although this differs from the default configuration of Boltz-2, Protenix-v1, and AlphaFold 3, it ensured that all models received the same sequence-only input. The absence of a consistent MSA benefit may reflect the short, repetitive Gly-X-Y structure of collagen mimetic peptides, which provides little sequence diversity for informative co-evolutionary constraints. Chai-1 retains its default ESM-2 embeddings and therefore differs in its internal sequence representation. All models were run using a single seed, following a five-seed sensitivity analysis (*Supplementary Information, S2*) that found negligible variation in lDDT and TM-score and limited variability in individual backbone RMSD, without altering the relative frequency of large-error predictions.

### 2.3 Evaluation

All predicted structures were generated in mmCIF format, matching the experimental reference structure files curated from the PDB. Predicted structures were evaluated using a common scoring pipeline implemented with Gemmi (v0.7.5) [40] for CIF parsing and atom/residue correspondence, and US-align (v20260527) [41] for structural alignment by TM-score maximization. Only heavy atoms were evaluated. Residue names were standardized before matching so that hydroxyproline was kept distinct from proline. Predicted and experimental residues were matched by sequence rather than by deposited chain identifiers or residue numbering. All possible assignments between the three predicted and three experimental chains were considered, and the assignment maximizing the number of matched residues was selected. Geometric agreement was used to resolve assignments with equal sequence coverage. This allowed equivalent homotrimer chains to be permuted and accommodated differences in chain naming and numbering among model outputs.

#### 2.3.1 Structure Accuracy Performance

Structural accuracy was evaluated using three complementary metrics: local Distance Difference Test (lDDT) [42], TM-score [43], and root-mean-square deviation (RMSD). Both all-atom and backbone lDDT and RMSD were calculated. TM-score was calculated using US-align in multimer mode (-mm 1), normalized by the experimental reference length, with RMSD calculated using the resulting global superposition. Together, lDDT measures local structural accuracy, while TM-score and RMSD provide complementary measures of global agreement.

All learned models were evaluated across the complete set of 80 structures. Direct paired comparisons with the deterministic pipeline were performed on the 75 structures successfully generated by all methods. Method coverage was reported separately so that the five deterministic-generation failures were not obscured by complete-case analysis. Aggregate performance was summarized using both the mean and median of each metric, together with per-structure paired comparisons. To assess potential training-set exposure, the benchmark was partitioned at each model’s stated training-data cutoff (Chai-1, 12 Jan 2021; AlphaFold 3 and Protenix-v1, 30 Sep 2021; Boltz-2, 1 Jun 2023), and performance evaluated separately on pre- and post-cutoff structures. CDSM requires no learned model training and uses a fixed empirical parameterization developed independently of the present benchmark. Sequence novelty itself is difficult to quantify directly because of collagen’s repetitive scaffold, limited representation of hydroxyproline in sequence-embedding methods, and the presence of three potentially dissimilar chains - each of different similarity - per structure. We therefore assessed novelty through the accumulation of newly deposited sequences and expansion of the observed Gly-X-Y triplet vocabulary after each cutoff.

#### 2.3.2 Computational Cost

Computational efficiency was evaluated using ten randomly selected structures from the benchmark dataset. For the deep-learning models, each structure was predicted using five seeds (seed = 1, 2, 3, 4, 5) to capture variation between runs. Wall-clock inference time and peak GPU memory were measured with predictions run sequentially on a dedicated NVIDIA L40S GPU via Modal. Initial model-loading time was recorded separately from subsequent “warm” predictions, with reported inference times representing the steady-state cost of generating an additional structure once model weights were loaded. CDSM was benchmarked on a single Apple M4 Pro CPU core, with OpenMM restricted to one thread, providing a conservative estimate for the parallelizable pipeline. Computational cost was calculated from median wall-clock time using Modal hardware prices (current as of August 2026) [44], with CDSM CPU time priced at Modal’s rate of $0.047 per core-hour. Peak memory measurements were used to identify the lowest-cost compatible GPU for each deep-learning model, on which costs were re-evaluated to estimate the minimum practical per-structure inference cost.

## 3 RESULTS

### 3.1 CDSM extends THeBuScr to a broader range of collagen sequences

We first compare sequence coverage of CDSM and native THeBuScr across all 80 experimental structures. THeBuScr constructs only 7 structures (8.8%), corresponding to equal-length chains beginning with glycine and containing complete, correctly registered Gly-X-Y triplets. Following CDSM preprocessing, 75 of 80 structures (93.8%) are constructed wholly (7) or partially (68) using the original Rainey-Goh parameterization, a 10.7-fold increase in coverage. The remaining five contain extended internal disruptions of Gly-X-Y periodicity that cannot be parameterized. Across the 75 structures, terminal helical-type extension is required for 19, Kabsch-based terminal extrapolation for 68, and chain re-registration for 25; side-chain reconstruction and relaxation are applied to all.

We evaluate backbone (bb) and all-atom (aa) structural accuracy after each stage of the CDSM pipeline to determine the effect of successive modifications on performance. To enable consistent all-atom scoring, unrelaxed side chains are added after each intermediate stage, while side-chain energy minimization is applied only to the final relaxed structures in the last step.

Across metrics, the successive CDSM stages address distinct limitations of the initial “core only” structures. Since THeBuScr is evaluated only on the seven compatible structures, its performance cannot be directly compared with CDSM across the broader 75-structure set. Following preprocessing, the CDSM “core only” structures represent all 75 parameterizable sequences but retain only 89.8% mean residue coverage due to trimmed termini, resulting in a median TM-score of 0.808. In contrast, backbone lDDT remains high (0.983) and backbone RMSD low (1.14 Å), outperforming even the final relaxed structures on these metrics. However, lDDT and RMSD are calculated only over matched regions and are therefore blind to the missing termini, artificially favoring the incomplete core structures. TM-score, normalized by the length of the complete experimental reference, penalizes this incompleteness and is correspondingly lower.

Reconstruction of the trimmed termini restores 100% sequence coverage across all 75 parameterizable structures (“full seq”), increasing median TM-score from 0.808 to 0.867. Local accuracy initially decreases, with backbone RMSD increasing from 1.135 to 1.560 Å, as reconstructed chains can remain axially misregistered relative to the experimental structure. Subsequent chain re-registration corrects this for the affected structures and increases median TM-score to 0.875 and reduces backbone RMSD to 1.270 Å. Among the 25 structures re-registered, backbone RMSD improves in 19, with a median change of −4.0 Å. This is visible in Fig. 5b as several poor-performing structures move into the main distribution following re-registration.

**Figure 5:**
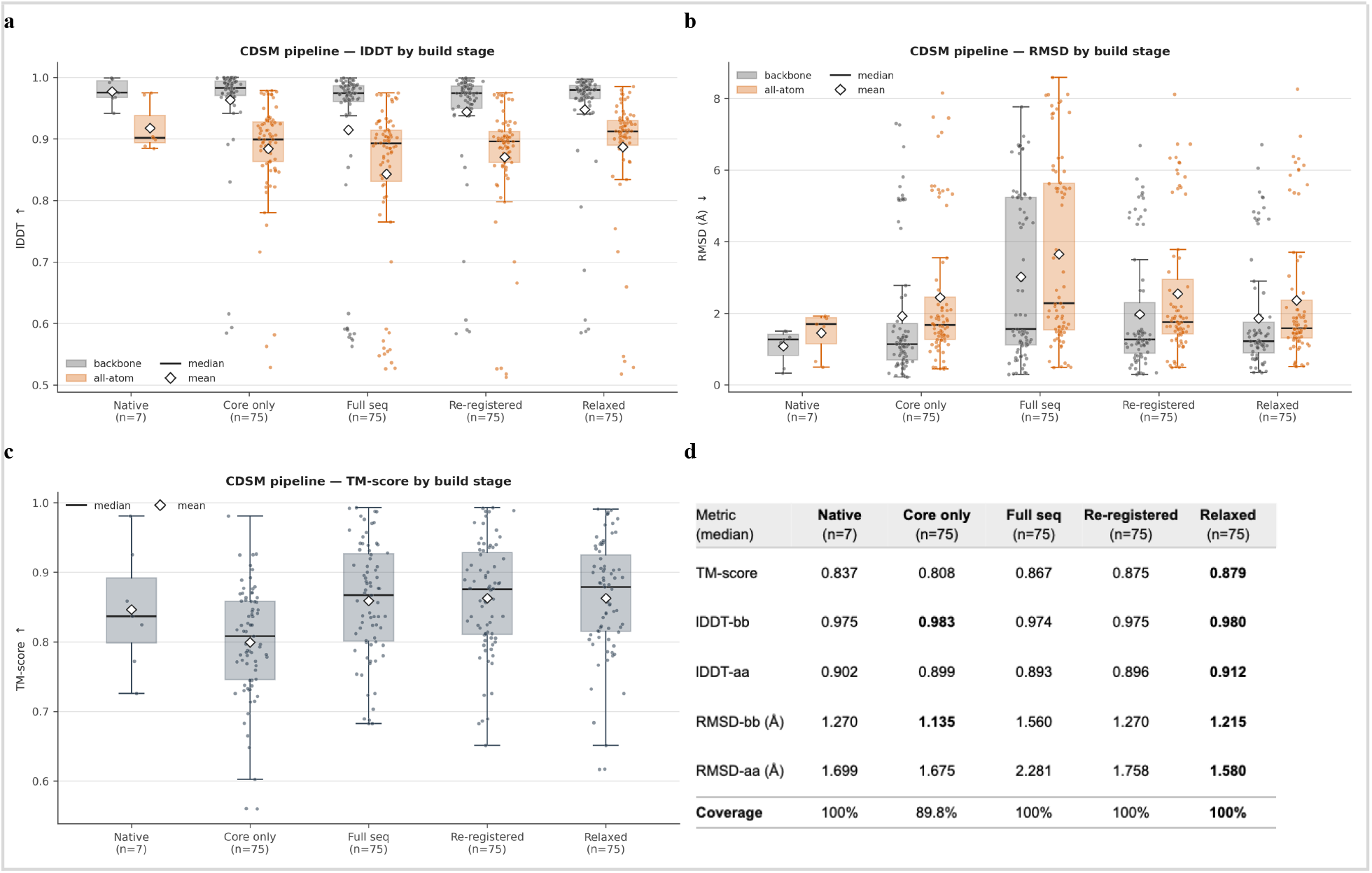
Evolution of CDSM structural accuracy through successive stages of reconstruction. Box plots show mean, median and IQR (whiskers = 1.5 IQR). Individual points represent structures. **a-b**. Backbone and all-atom lDDT and RMSD, respectively, across the CDSM pipeline. The initial “core only” structures retain 89.8% mean sequence coverage and therefore score favorably on lDDT and RMSD, which evaluate matched regions only. Restoring the complete sequence (“full seq”) initially worsens these metrics where terminal reconstruction introduces inter-chain misregistration; subsequent re-registration corrects the threading of 25 structures, recovering much of this accuracy, visible as poor-performing structures moving into the main distribution after this step. **c**. TM-score across build stages. Unlike lDDT and RMSD, TM-score penalizes the incomplete “core only” structures through normalization by the full reference length; restoring the termini therefore produces the largest increase, with smaller improvements following re-registration and relaxation. **d**. Median performance and sequence coverage at each stage. The final relaxed structures achieve the strongest overall combination of complete sequence coverage and structural accuracy, while the apparently superior backbone lDDT and RMSD of the “core only” structures reflect evaluation of incomplete chains. Native THeBuScr results are shown for the seven directly compatible structures but are not directly comparable with the broader 75-structure CDSM set.

**Figure 6:**
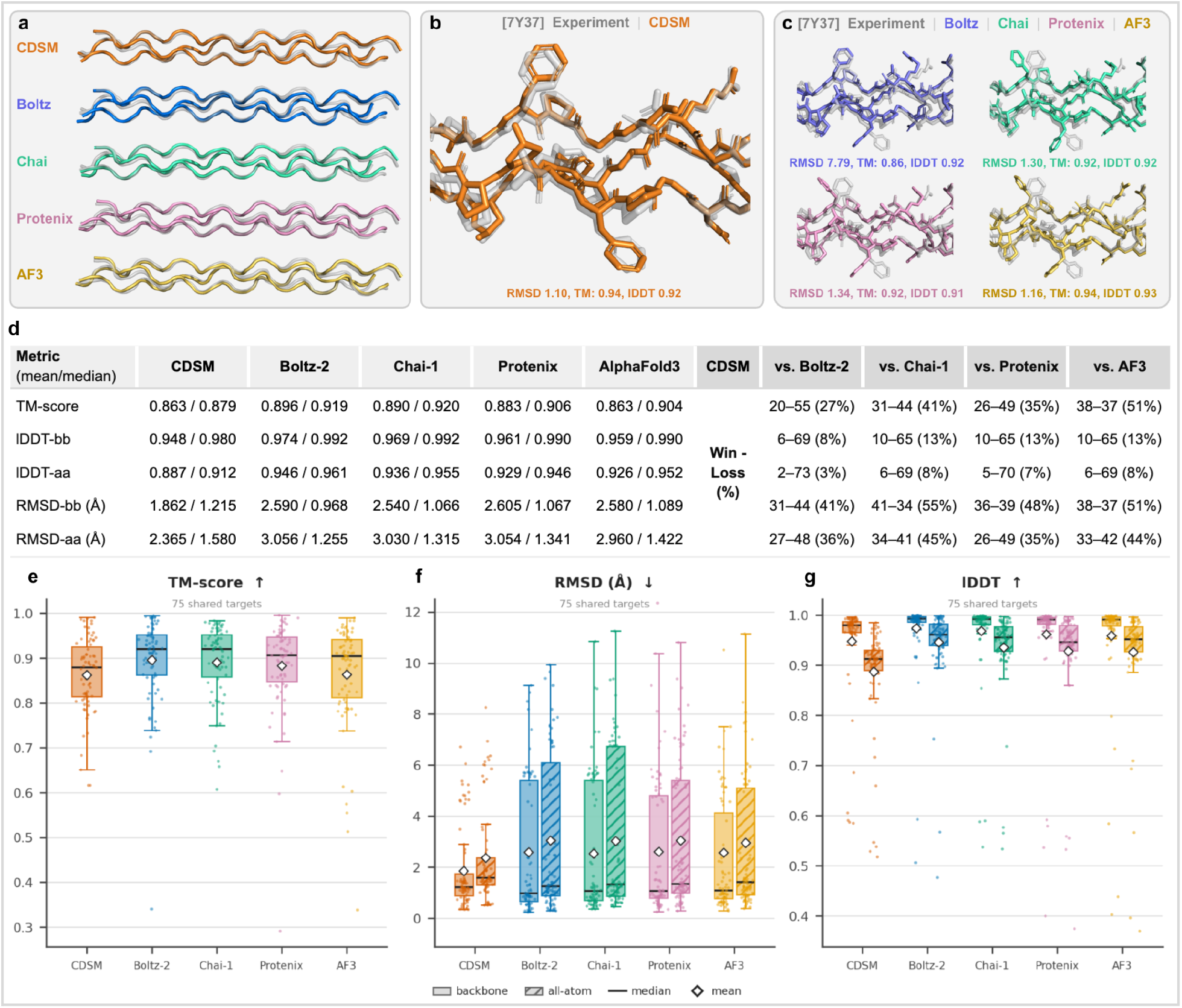
Structural accuracy of CDSM and deep-learning predictors across the 75 shared experimental structures. **a-c**. Representative overlays of predictions and the experimentally resolved structure of the complex heterotrimer 7Y37: (a) backbone comparison across all methods (b) detailed all-atom overlays for CDSM, and (c) the four deep-learning models. **d**. Summary of mean/median performance and pairwise CDSM win-loss counts across evaluation metrics. **e-g**. Distributions of TM-score, backbone and all-atom RMSD, and lDDT, respectively. CDSM achieves structural accuracy comparable to the learned models, with slightly lower TM-score and lDDT but competitive RMSD and fewer large-error predictions.

Finally, restrained energy minimization (“relaxed”) produces modest further improvements, increasing median TM-score from 0.875 to 0.879 and reducing backbone RMSD from 1.270 to 1.215 Å. Improvements are larger for side-chain-sensitive metrics, with all-atom lDDT increasing from 0.896 to 0.912 and RMSD decreasing from 1.758 to 1.580 Å. All-atom accuracy nevertheless remains lower than backbone accuracy, indicating that side-chain placement is a bigger source of error than the parameterized backbone geometry.

### 3.2 CDSM vs Generative AI: Comparison with learned structure predictors

Across all sequences, CDSM builds structures with accuracy comparable to predictions from the four evaluated structure prediction models, AlphaFold 3, Boltz-2, Chai-1, and Protenix-v1. While the four learned models generally achieve higher TM-score and lDDT scores, CDSM has the lowest mean backbone and all-atom RMSD (1.862 Å and 2.365 Å, respectively), despite slightly higher median RMSD. CDSM achieves a median TM-score of 0.879, compared with 0.906-0.920 for the learned models, and a median backbone lDDT of 0.980, compared with 0.990-0.992. The gap is wider for all-atom lDDT (0.912 for CDSM versus 0.946-0.961), indicating that much of the lDDT performance gap arises from side-chain rather than backbone geometry.

Although the learned models achieve lower median RMSD, CDSM exhibits fewer and less severe high-RMSD outliers from large-error predictions, resulting in lower mean RMSD. Thus, while the learned methods generally perform better in local structural similarity, CDSM achieves comparable global geometry with fewer large-RMSD predictions. At the individual-structure level, CDSM achieves broadly comparable backbone RMSD performance to AlphaFold 3 (51%), Chai-1 (55%), and Protenix-v1 (48%), with CDSM producing the lower-RMSD structure in approximately half of cases. This demonstrates competitive global accuracy across a substantial fraction of the benchmark. In contrast, CDSM achieves higher backbone lDDT on only 9-13% of structures, consistent with the learned models’ advantage in local, particularly side-chain, geometry.

### 3.3 CDSM trades modestly lower accuracy for robust global geometry

The difference in RMSD performance is most apparent in the tails of the distributions (Fig. 7a). Only five CDSM structures exceed 5 Å backbone RMSD, compared with 14 to 20 for the learned models, and its maximum error is 6.7 Å versus >10 Å for each learned model. The cumulative distributions cross at around 1.5 Å: below this CDSM resolves fewer targets than any of the models, above it more than all of them. CDSM is therefore modestly less accurate on typical structures but more robust to large global errors. Error is very position-dependent across all methods (Fig. 7b), with similar accuracy through the helix interior and most of the error concentrated at the termini. CDSM follows the same general profile but has a higher N-terminal error and an earlier error increase towards the C-terminus, although absolute errors at the termini remain comparable.

**Figure 7:**
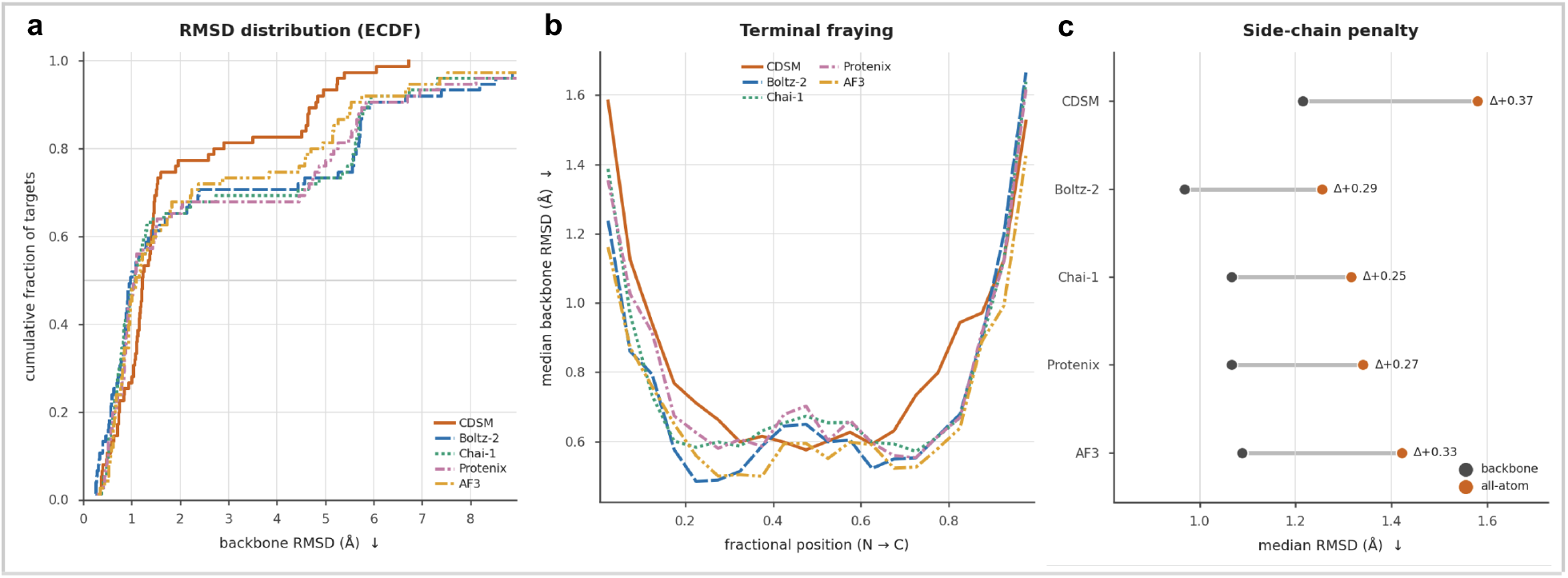
Error characteristics of CDSM and deep-learning structure predictors across the 75 shared targets. **a**. Empirical cumulative distribution of backbone RMSD, showing fewer large-error predictions for CDSM than Boltz-2, Chai-1, and Protenix-v1, and AlphaFold 3. CDSM’s worst case across all 75 targets is 6.7 Å against maxima above 10 Å for all four models. **b**. Median per-residue backbone RMSD along normalized chain position. All methods show low error throughout the triple-helical core and increased error toward the termini; CDSM exhibits greater N-terminal error and an earlier C-terminal increase, consistent with its geometric terminal reconstruction steps. **c**. Median backbone and all-atom RMSD, with Δ indicating the additional error associated with side-chain geometry. CDSM exhibits the largest backbone-to-all-atom penalty, 12-28% greater than those exhibited by the deep-learning models. All RMSD values are reported as a median throughout.

Moving from backbone to all-atom RMSD increases median error for every method, but the penalty is largest for CDSM (+0.37 Å vs +0.25-0.33 Å; Fig. 7c). This mirrors the all-atom lDDT deficit in Section 3.2 and indicates that CDSM’s main accuracy limitation lies in side-chain placement rather than in the construction of the backbone. Since CDSM side chains are constructed from force-field residue templates and energy-minimized, they produce locally minimized conformations, but not necessarily the specific conformers observed crystallographically.

Comparing by trimer type, CDSM achieves a median backbone RMSD of 1.10 Å on homotrimers, comparable to 0.89-1.15 Å for the learned models, but does less well on heterotrimers (1.37 Å versus 1.08-1.37 Å). Heterotrimers are more difficult across methods, although differences in lDDT and TM-score remain ≤0.01. Since many heterotrimers in this dataset also have unequal chain lengths and require terminal extensions and re-registration, their lower accuracy may partly reflect the additional reconstruction required. Together with the previous results, this suggests that CDSM captures the triple-helical core well, and that its errors emerge from terminal reconstruction, chain register, and side-chain placement.

### 3.4 CDSM Becomes More Competitive Relative to Learned Models on Post-Training-Cutoff Structures

To assess whether the learned models benefit from exposure to pre-cutoff structural data, we separately evaluate performance before and after each model’s stated training-data cutoff. Most benchmark structures predate the respective cutoffs (59 to 75%, depending on model) and therefore represent possible training-set exposure. CDSM, which has no learned training set and whose fixed empirical parameterization predates the training cutoffs of all four learned models, is evaluated on the same subsets to assess comparative performance on post-cutoff data to contextualize differences in structural difficulty. The post-cutoff period also coincides with substantial expansion of the experimentally observed Gly-X-Y triplet vocabulary (Fig. 8a), indicating that newer structures introduce sequence contexts not seen in earlier benchmark structures.

**Figure 8:**
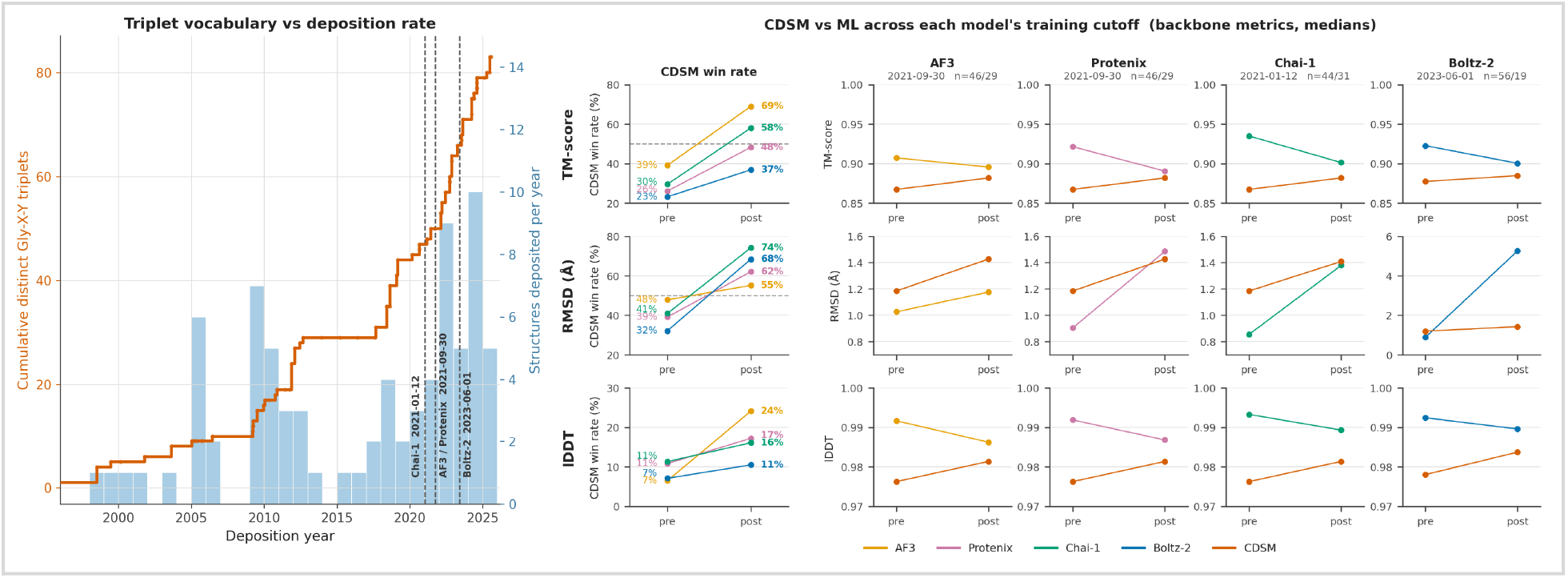
Dataset novelty and CDSM performance relative to learned models across training-data cutoffs. **a**. Growth of the experimentally observed Gly-X-Y triplet vocabulary and new annual structure deposition across the benchmark; dashed lines show the stated training-data cutoffs. Triplet vocabulary increased from 50 distinct triplets at the AlphaFold 3/Protenix-v1 cutoff (2021) to 83 in the complete dataset, indicating substantial sequence novelty in the post-cutoff data. **b**. CDSM pairwise win rates and median backbone TM-score, RMSD, and lDDT before and after each model’s respective cutoff. CDSM becomes more competitive across all metrics post-cutoff as learned-model performance generally moves toward or below CDSM performance; all-atom metrics show the same overall trend. Boltz-2 shows the largest shift in backbone RMSD (0.89 to 5.26 Å), with its 19 post-cutoff predictions separating into nine low-error (0.27-2.39 Å) and ten high-error (5.26-6.65 Å) structures, a result of discrete global errors, mostly from chain misregistration.

Across all cutoffs, CDSM improves relative to the learned models on post-cutoff structures (Fig. 8b), with its pairwise win rate increasing across all five metrics. Aggregate CDSM win rates increase from 29% to 55% for TM-score and 40% to 65% for backbone RMSD. The increase is smaller for lDDT (7% to 18%), where CDSM continues to underperform the learned methods. As CDSM performance changes comparatively little across the same subsets, the relative shift is driven primarily by degradation of the learned models. This pattern is consistent with, but does not establish, a contribution from pre-cutoff training-data exposure, because post-cutoff structures may also differ systematically in sequence composition and structural difficulty.

### 3.5 CDSM structural limitation: Unable to handle internal triplet interruptions

CDSM fails to build 5 of the 80 experimental structures: 1EI8, 6M80, 5K86, 7LXQ, 7LXP. In each case, prolonged or repeated interruptions to the Gly-X-Y periodicity leave THeBuScr’s 5-triplet propensity-averaging window without a valid Gly-led triplet at some point along the chain, causing a divide-by-zero error during helix-type assignment. Because CDSM’s frame correction operates as a single global reframe, it cannot resolve a break occurring partway through a chain; doing so would require locally approximating melting temperature or helical propensity around the interruption. This is a structural limitation of the current pipeline rather than a tunable parameter, and is a concrete target for future work.

By contrast, all four deep-learning models predict these five structures, representing a clear coverage advantage. However, all models find these structures markedly harder than the shared 75-structure set: median TM-score falls from 0.91 to 0.63, backbone and all-atom lDDT from 0.99 to 0.90 and 0.95 to 0.85, respectively, while backbone and all-atom RMSD increase from 1.07 to 4.92 Å and 1.32 to 5.94 Å. The much larger degradation in RMSD than lDDT suggests that local triple-helical geometry is partly retained while global structural arrangement is disrupted. The same Gly-X-Y interruptions that prevent CDSM construction also present a substantial challenge to the learned models, indicating that non-canonical collagen sequences remain difficult across methods.

### 3.6 Computational cost

CDSM is substantially faster and less expensive than all four deep-learning methods (*Table 2*). In the standardized comparison, with all deep-learning models run on the same NVIDIA L40S GPU, median prediction times ranged from 31.9-78.9 s per structure, compared with 2.4 s for CDSM on a single CPU core. This corresponds to $0.017-0.043 per structure for the deep-learning models versus $3.1 × 10^·5^ ($0.000031) for CDSM, making CDSM approximately 560-1,380× less expensive. At these rates, one US dollar generates approximately 32,000 CDSM structures, compared with 23 to 59 deep-learning predictions.

This comparison is conservative with respect to deep-learning deployment because the L40S provides substantially more memory than required by several models. We therefore repeat inference using the lowest-cost compatible GPU available, an NVIDIA L4. Despite 1.39-2.03× longer inference times, its lower hourly cost reduces deep-learning prediction costs by 16-43%, to $0.013-0.025 per structure, with the smallest saving for Chai-1 due to its larger 2.03× slowdown. However, even under this hardware-optimized comparison, CDSM remained 400-790× less expensive than the deep-learning models.

Unlike CDSM, the deep-learning models additionally incur substantial one-time model-loading overhead when initialized in a fresh container, ranging from 491 s for Boltz-2 to 1,030 s for Chai-1 (568 s for Protenix-v1 and AlphaFold 3 incurred no comparable overhead as parameters were pre-staged on persistent storage); these cold-start costs were excluded from the steady-state comparisons above. As an external reference, the measured self-hosted Boltz-2 cost ($0.017 per structure on L40S) is within 1.5× of the commercially hosted Boltz-2 API price ($0.025 per structure), supporting the magnitude of the cost estimates.

## 4 DISCUSSION

### 4.1 A compact geometric representation efficiently compresses the collagen structure-prediction problem

This work finds no universally superior method for collagen structure prediction but shows that a compact, collagen-specific geometric representation can approach the accuracy of frontier deep-learning models at substantially lower computational cost. CDSM is particularly competitive in backbone and global geometry and produces fewer large errors, while the learned models retain an advantage in local and side-chain accuracy. At 400 to 790× lower computational cost, these results raise the broader question of geometric representation as simple as CDSM’s can approach the performance of substantially more complex learned models.

The findings illustrate the value of embedding domain-specific constraints into an analytical representation of a structure-prediction problem. Collagen occupies a highly constrained structural manifold defined by Gly-X-Y periodicity, a narrow range of backbone conformations, and its conserved triple-helical topology and chain stagger. Through the Rainey-Goh parameterization, CDSM encodes these constraints, substantially reducing the structural space that must be explored. The results further suggest that much of the representational capacity of general-purpose generative models may be unnecessary for structurally constrained problems: once relevant structural rules are known, a relatively simple model can capture much of the same geometry. This constraint may also contribute to CDSM’s robustness by limiting the range of possible global errors, consistent with its maximum backbone RMSD of 6.7 Å compared with predictions exceeding 10 Å for each learned model.

The training-cutoff analysis provides further evidence of the robustness of this analytical approach relative to learned models. While the learned models all degrade in performance on post-cutoff structures, CDSM maintains broadly similar accuracy across the benchmark, resulting in its aggregate win rates increasing substantially across all evaluation metrics. Although this analysis does not demonstrate memorization or confirm exposure to individual structures, it is consistent with some benefit from pre-cutoff training data and suggests that CDSM’s explicit, fixed empirical representation generalizes comparatively well across structures.

### 4.2 Implications for high-throughput collagen design

The practical importance of the remaining accuracy gap depends on how the structures are used. CDSM was developed in part to enable high-throughput prediction of collagen structures for data-driven design and downstream mechanical simulation, where correct triple-helical topology and chain register may be more important than small differences in local atomic coordinates. CDSM’s backbone errors are comparable in magnitude to the approximately 1 to 2 Å fluctuations for equilibrated collagen triple helices [45], suggesting that sub-Ångström differences in starting coordinates may be less consequential after equilibration. This remains to be tested directly by comparing the mechanical properties predicted from structures generated by different methods.

For high-throughput applications, computational cost and robustness may therefore be as important as marginal improvements in accuracy. Large errors can propagate unnoticed through automated pipelines, while small local deviations may be inconsequential after equilibration. CDSM’s comparatively fewer large errors and large cost advantage make it suitable for structural screening, where large candidate sets can be generated efficiently, reproducibly without model training or specialized accelerator hardware. CDSM therefore offers an alternative when the objective is reliable, high-throughput structure generation with reproducibility, interpretability, and robust global geometry, rather than maximum local atomic accuracy.

### 4.3 Limits of a constrained geometric representation

The same constrained geometric representation that enables CDSM’s efficiency and performance also introduces limitations. In its current form, CDSM cannot construct extended internal disruptions of Gly-X-Y periodicity, which limits its coverage to 75 of the 80 collagen sequences in the benchmark. Notably, these five structures are also much more difficult for the learned models, whose median backbone RMSD increases from 1.07 to 4.92 Å and TM-score decreases from 0.91 to 0.63. Nevertheless, these failures illustrate a broader limitation of compact representations: while a strong structural prior performs remarkably well within the structural space it describes, it becomes less effective as sequences depart from the rules it encodes. It is here that the flexibility of general-purpose learned models becomes valuable; in this case allowing them to represent structures beyond the assumptions built into CDSM.

Side-chain accuracy represents a further distinct limitation. CDSM’s empirical representation primarily determines backbone geometry, while side chains are placed using force-field templates and restrained minimization. The resulting structures reproduce experimental side-chain coordinates less accurately than the learned methods since CDSM’s representation does not explicitly capture environmental effects such as solvent and crystal packing that can influence experimentally observed side-chain conformations. CDSM may therefore be less suitable for applications requiring precise local interaction geometry.

The benchmark itself also imposes limitations. The experimental dataset is relatively small and contains inherent sequence similarity, reflecting collagen’s conserved and repetitive sequence architecture. It is deliberately restricted to the 20 standard amino acids and 4-hydroxyproline (HYP), capturing the predominant chemistry of natural collagen while excluding rare post-translational variants and synthetic amino-acid analogues. The pre/post-cutoff analysis should also be interpreted cautiously: deposition dates do not establish whether individual structures were included in model training, and collagen’s repetitive sequences make novelty difficult to quantify, although continued growth in the observed Gly-X-Y triplet vocabulary indicates the introduction of new sequence contexts after the model cutoffs. Finally, experimental structures are references rather than perfect ground truth, particularly for flexible termini and solvent-exposed side chains whose conformations may be uncertain or influenced by the crystallographic environment. Differences from deposited coordinates therefore do not necessarily correspond directly to physical error.

### 4.4 Towards autonomously discovered representations

CDSM provides an example of a compact scientific representation derived manually from known structural constraints and a relatively small empirical parameterization. Its performance points to a broader opportunity to determine whether similarly efficient representations can be discovered for other proteins or structural domains. Collagen’s unusually constrained geometry makes such a representation relatively straightforward to derive manually; for more complex protein structures, identifying the appropriate rules and parameterizations may exceed what can readily be formulated by hand.

Emerging AI-led approaches provide a possible route to automate such a search, especially for open-ended problems for which clear success metrics exist [33]. Other, related methods such as AlphaTensor and FunSearch have also demonstrated that generative AI can be used to search directly over algorithms or executable functions to discover solutions that improve upon established human-designed approaches [46, 47]. A similar principle as what was done in [33] to identify a model to predict protein B-factors could be applied to protein structures: candidate geometric rules, structural representations, and empirical parameterizations could be generated and evaluated against experimental structures for accuracy, generalization, complexity, and computational cost. Self-revising discovery systems could further allow these representations to be continually challenged, reparametrized, and refined as new structural data become available [33].

Such an approach would use the capability of deep learning and more specifically generative AI methods differently from current end-to-end structure predictors: rather than learning a complex and general mapping from sequence to structure, models would search for compact algorithms that capture the underlying structural rules themselves. CDSM demonstrates the potential value of this approach when those rules can be identified explicitly. Whether similarly compact representations exist for less constrained protein domains remains an open question, but their discovery could provide interpretable and computationally efficient alternatives to today’s learned structure predictors.

### 4.5 Further work

Several immediate extensions could improve CDSM. The original Rainey-Goh parameterization, derived from only ten available structures, could now be re-derived using the substantially larger collagen structural corpus, although improvements would primarily affect the backbone, which already performs well. More importantly, the expanded dataset may enable better treatment of extended Gly-X-Y periodicity disruptions. Possible approaches include extending the existing propensity parameterization to previously unseen sequence contexts, directly parameterizing the local geometry of common interruption types, or reconstructing interrupted regions between parameterized triple-helical segments using geometric loop-closure methods or the same Kabsch extrapolation used here for terminal residue coordinates. Side-chain accuracy may also be improved through alternative force-field templates or explicit rotamer sampling. Full-dataset evaluation of the learned methods under their default MSA configurations would further strengthen the comparison, although preliminary tests found little difference for these short, repetitive sequences.

Importantly, the speed and low computational cost of CDSM make it attractive for structure prediction and design screening at substantially larger scales. Ongoing work within the LAMM group at MIT is using predicted structures for high-throughput mechanical simulation and candidate-sequence screening, as well as graph neural network approaches to learn local and global sequence-structure-property relationships. Future work to compare the mechanical properties after molecular dynamics equilibration and simulation on structures generated by CDSM and the learned models could establish whether the small differences in coordinate accuracy observed here translate into meaningful differences for downstream collagen design.

## 5 CONCLUSION

This work demonstrates that explicitly encoding collagen’s strong geometric constraints into an efficient algorithm can reproduce experimental triple-helical structures with accuracy approaching frontier deep-learning predictors while requiring orders of magnitude less computation and producing fewer severe global errors. Learned models remain preferable where maximum local and side-chain accuracy is required, whereas CDSM provides a reproducible, interpretable, and computationally lightweight approach well suited to high-throughput generation of collagen triple-helical structures for downstream simulation and design. More broadly, these results show that large representational capacity need not always translate into proportionally greater predictive value: when the relevant rules of a scientific problem can be found and encoded, the right representation can dramatically compress the problem while keeping most of the predictive accuracy.

## AUTHOR CONTRIBUTIONS

The manuscript was written with contributions from all authors. All authors have given their approval to the final version of the manuscript. Conceptualization, methodology, and computational analysis: B.H. and M.B.; writing, original draft: B.H. and M.B; writing, reviewing and editing: M.J.B, B.H, M.B; supervision: M.J.B. All authors have reviewed and given approval to the final version of the manuscript.

## SUPPORTING INFORMATION

Additional details are provided on experimental structure dataset curation and processing (S1); and supplementary structure prediction results and analyses (S2).

## DATA AVAILABILITY

The curated experimental dataset, predicted structures, and data underlying the analyses presented in this study are openly available at: https://huggingface.co/datasets/lamm-mit/collagen-cdsm_benchmarking_data.

Code used for dataset curation, structure prediction, evaluation, and analysis is openly available at https://github.com/lamm-mit/Collagen-Structure-Modeler.

## CONFLICTS OF INTEREST

The authors declare no conflicts of interests.

## ACKNOWLEDGEMENTS

Support from MGAIC is acknowledged.

## SUPPLEMENTARY INFORMATION

### S1. Dataset: Experimental Structure Dataset Curation

Experimental collagen triple-helix structures were retrieved from the RCSB PDB on 2026-07-01 using the RCSB Search API (v2). The initial query required entries to match the keywords “collagen” or “triple helix”, contain a polymer chain count that is a positive multiple of three (to capture asymmetric units comprising one or more triple helices), and consist exclusively of protein chains. This returned 758 candidate structures. Structures were then filtered sequentially as follows, with each structure rejected at the first failed criterion:

#### Notes on filter design

The glycine content filter (≥25%) is applied first among post-download filters because it is the most discriminating criterion: non-collagen trimeric proteins captured by the keyword search, including coiled-coils, trimeric antibodies, and viral coat proteins, typically exhibit substantially lower glycine fractions. The theoretical Gly fraction of a canonical Gly-X-Y repeating collagen structure is roughly one third; the 25% threshold accommodates internal disruptions and chain termini that may not align with a complete triplet. The residue allow list is limited to the 20 standard amino acids plus (4R)-hydroxyproline, HYP. This removes 15 structures from the dataset. Two structures containing close hydroxyproline analogues, (4S)-hydroxyproline (HZP) in 3B2C and 3-hydroxyproline (HY3) in 2G66, are excluded despite being genuine collagen post-translational modifications; this represents a limitation of the current benchmark. The remaining 13 structures removed at this step contain other non-allowed residues, predominantly synthetic analogues, including one (3DMW) containing selenomethionine, which could be recovered by treating it as methionine. The chain-length bounds (10–200 residues) capture the typical range of collagen mimetic peptides while excluding very short fragments and longer collagenous proteins outside the benchmark scope.

#### Chain count and multi-copy handling

The API query accepts chain counts of 3, 6, 9, or 12 to recover structures where the asymmetric unit contains more than one triple helix, a common crystallographic outcome. For such structures, chains were grouped into triple helices by spatial distance, with chains connected when their minimum Cα-Cα distance was <5.5 Å and connected components of exactly three chains identified as individual helices. For structures containing multiple helices, one representative helix was retained: the component containing the earliest chain in ATOM-record order. Sequence extraction used ATOM records exclusively, with hydroxyproline represented internally as O.

#### Reproducibility

The full pipeline is implemented in download_collagen.py and archived alongside the dataset on Hugging Face. Minor variation in candidate counts between runs may occur as the PDB index is updated.

**Table S.1:** Sequential filtering of RCSB PDB structures used to construct the experimental collagen benchmark dataset.

| Filter | Criterion | Removed | Remaining |
| --- | --- | --- | --- |
| Initial API query | Keyword + chain count $\in \{3,6,9,12\}$ + protein only, retrieved July-2026 | — | 758 |
| Glycine content | Gly fraction of ATOM sequence $< 25\%$ | 651 | 107 |
| Residue allow list | Any chain contains residues outside the 21-type AAs (20 standard + HYP) | 15 | 92 |
| Structural continuity | Internal residue numbering gap $> 5$ within any chain | 1 | 91 |
| Chain length | Any chain $< 10$ or $> 200$ residues | 7 | 84 |
| Deduplication | 9I99 / ~~9I9A~~ / ~~9IBU~~: identical (PPG) □ entries; 6VZX / ~~3T4F~~ & 5Y45 / ~~5Y46~~: duplicate. | 4 | 80 |

### S2. Seed Sensitivity Analysis for Single-Seed Accuracy Benchmark

The accuracy benchmark compares one diffusion sample per learned model (*Table 1*). Because these models are stochastic, we evaluate the sensitivity of the accuracy results to seed choice. The seed sweep used the same compute-cost benchmark set (*Section 2.3.2*): ten benchmark structures spanning the range of sequence lengths were evaluated over five seeds for each learned model. Predictions were scored using the same metrics and experimental references as the main benchmark. Since CDSM is deterministic, it was evaluated just once per target.

**Table 1:**
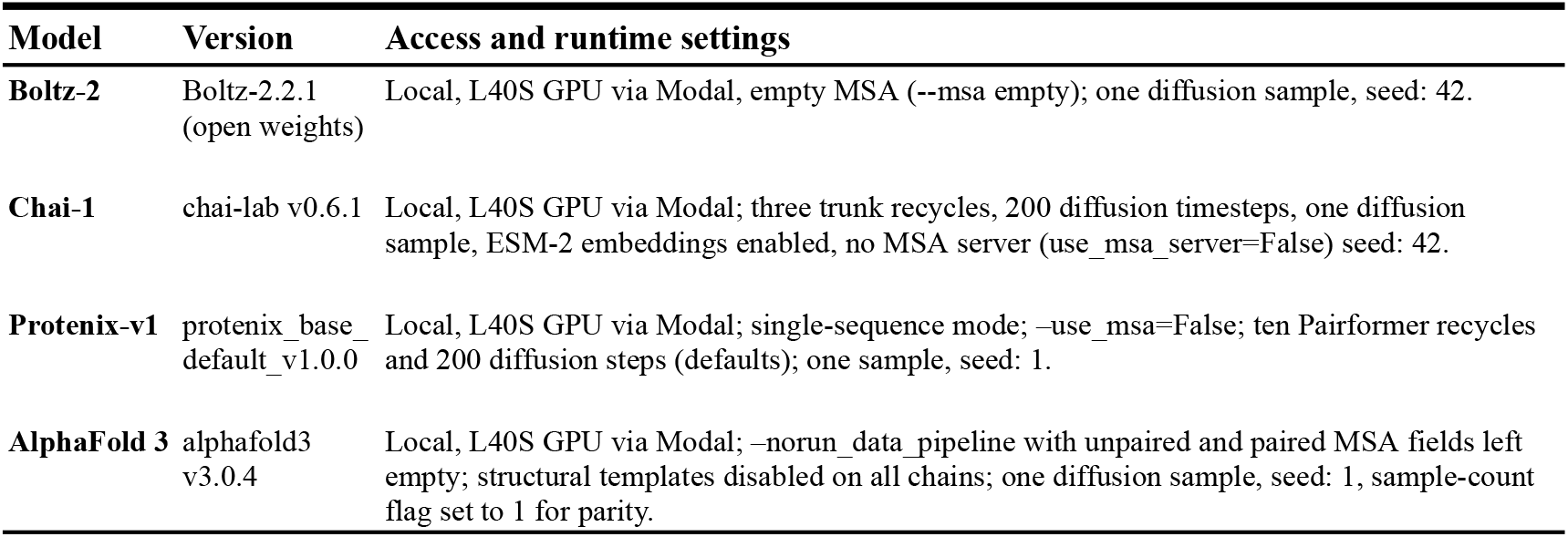
Access and runtime settings for each deep-learning model were mostly chosen as default and kept equivalent between models where supported. All models were run in single-sequence mode without MSA, after a preliminary comparison yielded no substantial difference. Single-seed predictions are justified by a seed-sensitivity analysis, detailed in S2.

| Model | Version | Access and runtime settings |
| --- | --- | --- |
| **Boltz-2** | Boltz-2.2.1<br>(open weights) | Local, L40S GPU via Modal, empty MSA (--msa empty); one diffusion sample, seed: 42. |
| **Chai-1** | chai-lab v0.6.1 | Local, L40S GPU via Modal; three trunk recycles, 200 diffusion timesteps, one diffusion sample, ESM-2 embeddings enabled, no MSA server (use_msa_server=False) seed: 42. |
| **Protenix-v1** | protenix_base_<br>default_v1.0.0 | Local, L40S GPU via Modal; single-sequence mode; --use_msa=False; ten Pairformer recycles and 200 diffusion steps (defaults); one sample, seed: 1. |
| **AlphaFold 3** | alphafold3<br>v3.0.4 | Local, L40S GPU via Modal; --norun_data_pipeline with unpaired and paired MSA fields left empty; structural templates disabled on all chains; one diffusion sample, seed: 1, sample-count flag set to 1 for parity. |

**Table 2:** Computational efficiency and per-structure cost of CDSM and deep-learning structure predictors. Deep-learning models were benchmarked on a standardized NVIDIA L40S and the lowest-cost compatible GPU (NVIDIA L4). Each model was given 10 structures × 5 diffusion seeds per GPU with execution order randomized to mitigate seed confounding with run position. Times report median steady-state inference, excluding model loading; ± is the standard deviation across per-seed medians. Absolute runtimes on shared cloud infrastructures varied by 21-56% between independent measurement sessions, thus per-structure costs are order-of-magnitude estimates. Costs use Modal prices from 12 August 2026. CDSM is deterministic and has no seed dependence and was benchmarked on a single Apple M4 Pro CPU core over the same structures.

| Method | Hardware | Time (s) | Peak GPU memory | Cost/structure | vs. CDSM |
| --- | --- | --- | --- | --- | --- |
| **CDSM** | M4 Pro CPU | 2.4 | — | \$0.000031 | — |
| **Boltz-2** | L40S / L4 | 31.9 ± 0.2 / 55.9 ± 0.5 | 3.48 GiB | \$0.017 / \$0.013 | 560 / 400× |
| **Chai-1** | L40S / L4 | 39.9 ± 0.2 / 81.1 ± 0.2 | 14.69 GiB | \$0.022 / \$0.018 | 700 / 590× |
| **AlphaFold 3** | L40S / L4 | 57.4 ± 0.4 / 87.6 ± 1.1 | 5.31 GiB | \$0.031 / \$0.020 | 1,010 / 630× |
| **Protenix-v1** | L40S / L4 | 78.9 ± 1.3 / 110.0 ± 1.3 | 4.56 GiB | \$0.043 / \$0.025 | 1,380 / 790× |

As shown in *Table S.2*, seed sensitivity is small for local metrics but greater for backbone RMSD. The median per-structure standard deviation across seeds is 0.015-0.025 for TM-score and ≤0.004 for lDDT across the four learned models. Backbone RMSD variability arises primarily from chain misregistration, resulting in errors of 4-8 Å in a small number of structures.

Overall, the learned models achieve higher lDDT and TM-score than CDSM, with little sensitivity to seed choice. In contrast, backbone RMSD is more seed-sensitive: median rankings and individual-structure comparisons vary across seeds. However, CDSM produces no backbone-RMSD errors above 5 Å in this subset, whereas each learned model produces at least one across every seed tested. Accordingly, RMSD median rankings and per-structure win rates in the main benchmark should be interpreted as approximate, while the overall comparison of large-RMSD errors is robust to seed choice.

**Table S.2:**
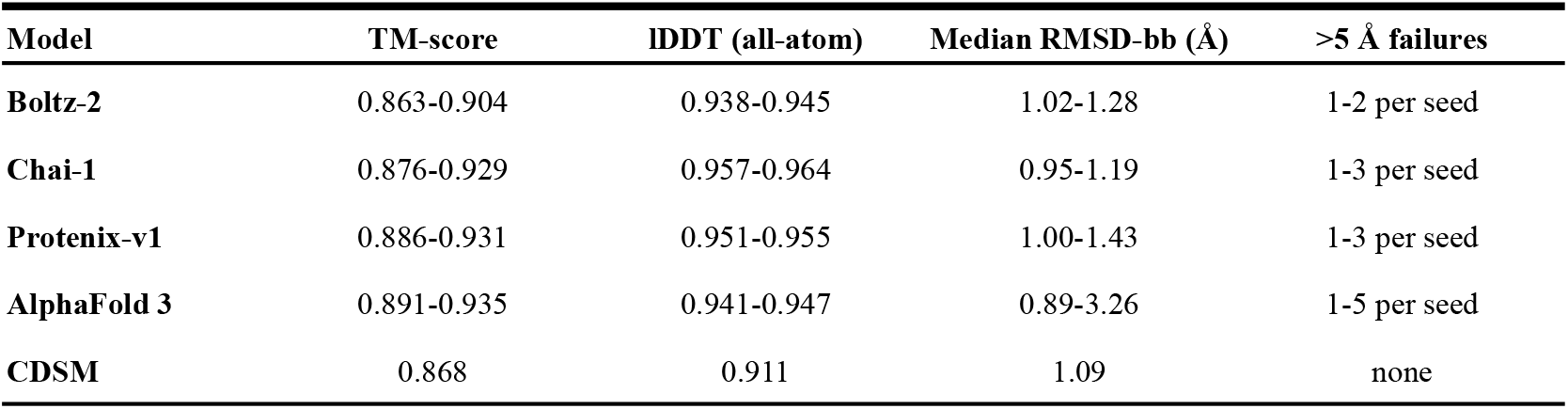
Per-seed medians across ten structures (ranges over seeds 1-5, L40S GPU) and occurrences of predictions with backbone RMSD >5 Å across the five seeds per target.

## Notes

https://github.com/lamm-mit/Collagen-Structure-Modeler

https://huggingface.co/datasets/lamm-mit/collagen-cdsm_benchmarking_data

### Competing Interest Statement

The authors have declared no competing interest.

### Funding

- Massachusetts Institute of Technology: MGAIC

